# No effect of monetary reward in a visual working memory task

**DOI:** 10.1101/767343

**Authors:** Ronald van den Berg, Qijia Zou, Yuhang Li, Wei Ji Ma

## Abstract

Previous work has shown that humans distribute their visual working memory (VWM) resources flexibly across items: the higher the importance of an item, the better it is remembered. A related, but much less studied question is whether people also have control over the *total* amount of VWM resource allocated to a task. Here, we approach this question by testing whether increasing monetary incentives results in better overall VWM performance. In three experiments, subjects performed a delayed-estimation task on the Amazon Turk platform. In the first two experiments, four groups of subjects received a bonus payment based on their performance, with the maximum bonus ranging from $0 to $10 between groups. We found no effect of the amount of bonus on intrinsic motivation or on VWM performance in either experiment. In the third experiment, reward was manipulated on a trial-by-trial basis using a within-subjects design. Again, no evidence was found that VWM performance depended on the magnitude of potential reward. These results suggest that encoding quality in visual working memory is insensitive to monetary reward, which has implications for resource-rational theories of VWM.

## INTRODUCTION

A central question in research on human visual working memory (VWM) is how much flexibility exists in how the system distributes its resource across encoded items (Luck & Vogel, 2013; Ma et al., 2014). The answer to this question partly depends on how one conceptualizes the nature of VWM resource. One class of models postulates that VWM consists of a small number of “slots” that each provide an indivisible amount of encoding resource (e.g., (Awh et al., 2007; Cowan, 2001; Luck & Vogel, 1997; Rouder et al., 2008; Zhang & Luck, 2008)). Since the number of slots is typically assumed to be very small (3 or 4), these models allow for virtually no flexibility in resource allocation. A competing class of models conceptualizes VWM as a continuous resource (e.g., (Bays & Husain, 2008; Fougnie et al., 2012; Keshvari et al., 2013; Shaw, 1980; van den Berg et al., 2012; Wilken & Ma, 2004)), sometimes in combination with a limit on the number of encoded items (Sims et al., 2012; van den Berg et al., 2014). Since a continuous resource can be divided into arbitrarily small packages, these models allow for a high degree of flexibility in resource allocation.

Several recent studies have found evidence for flexibility in VWM resource allocation. First, it has been found in multiple experiments that when one item in a stimulus array is more likely to be selected for test than other items (i.e., have a higher “probing probability”), subjects remember this item with better precision (Bays, 2014; Bays et al., 2011; Emrich et al., 2017; Gorgoraptis et al., 2011; Yoo et al., 2018; Zokaei et al., 2011). Similarly, people remember items associated with a higher reward better than items associated with a lower reward (Klyszejko et al., 2014). In addition, it has been reported that subjects can make a tradeoff between the number of items in VWM and the quality with which they are encoded (Fougnie, Cormiea, Kanabar, & Alvarez, 2016; however see Zhang & Luck, 2011). By allocating more resources to the more important items within a display, subjects in these studies increased their performance compared to what it would have been if they had encoded all items within each display with the same precision. This suggests that VWM resource allocation may be driven by a rational policy.

We recently formalized this suggestion by modeling VWM as a rational system that balances the amount of invested resource against expected task performance: the more there is at stake, the more resource is allocated for encoding (van den Berg & Ma, 2018). This “resource-rational” interpretation of VWM predicts two kinds of flexibility in the allocation of VWM resource. First, items of unequal importance are assigned unequal amounts of encoding resource. For example, when one item in a memory array is more likely to be probed, the resource-rational strategy would be to encode it with higher precision than the other items. In support of this prediction, we found that the model provided excellent quantitative fits to data from previous experiments that varied probing probabilities (Bays, 2014; Emrich et al., 2017). A second prediction made by the model is that *tasks* of unequal importance are assigned unequal amounts of *total* resource: the higher the incentive to perform well on a task, the more VWM resource a subject should be willing to invest. In support of the second kind of flexibility, it has been found that subjects who are encouraged to “try to remember all items” in a change detection task have higher estimated numbers of slots than subjects who are told to “just do your best” or to “focus on a subset” (Bengson & Luck, 2016). Moreover, there is evidence that people can flexibly trade off resources between auditory and visual working memory based on the amount of reward associated with each task (Morey et al., 2011). Finally, there are indications that retro-cueing can increase net VWM capacity (Myers et al., 2018).

In the present study, we examine whether the precision with which people encode a set of stimuli in VWM depends on the amount of monetary reward they get for good performance. We performed three experiments in which subjects earned a performance-contingent monetary bonus on top of a base payment. When encoding is costly, a rational observer should adjust its total amount of invested VWM resource to the amount of performance-contingent bonus: the higher the potential bonus, the more effort should be put into the task. We did not find evidence for such an effect in any of the experiments. In opposition to the prediction following from a resource-rational theory of VWM (Van den Berg & Ma, 2018), the present results suggests that encoding precision in VWM is insensitive to monetary reward.

## EXPERIMENT 1

### Data and code availability

All data, Matlab analysis scripts to reproduce figures of results, and JASP files with statistical analyses are available at https://osf.io/mwz27/.

### Recruitment

Subjects were recruited on the Amazon Mechanical Turk platform, where the experiment was posted as a “Human Intelligence Task”. The experiment was visible only to subjects who were located in the USA, had not participated in the experiment before, and had an approval rate of 95% or higher. A total of 355 subjects signed up, of which 156 were disqualified due to failing the post-instruction quiz (see below). The remaining 199 subjects were randomly assigned to four groups (*n*=49, 47, 47, 46) that differed in the total amount of bonus they could earn by performing well ($0, $2, $6, $10). Besides the bonus, subjects received a $1 base payment. The experiment was approved by the Institutional Review Board of New York University.

### Stimuli and task

On each trial, the subject was presented with 1, 2, 4, 6, or 8 Gabor patches, which were placed along an invisible circle around a central fixation point (Figure 1A). We refer to the number of presented items as the set size, which varied from trial to trial in a pseudo-random manner. The orientation of each patch was drawn independently from a uniform distribution over all possible orientations. The stimulus appeared for 50 milliseconds and was followed by an empty screen with a duration of 1 second (memory period). Thereafter, a randomly oriented Gabor patch appeared at one of the previous stimulus locations, whose initial orientation was randomly drawn and could be adjusted through mouse movement. The task was to match the orientation of this probe stimulus with the remembered orientation at that location. Only one item was probed on each trial. After submitting the response, the error between the correct orientation and the reported orientation, *ε*, was converted into an integer score between 0 and 10, with more points assigned for smaller errors (see Appendix for a visualization of the scoring function). Feedback was provided after each trial by showing the obtained score and two lines that corresponded with the correct and responded orientations.

**Figure 1.**
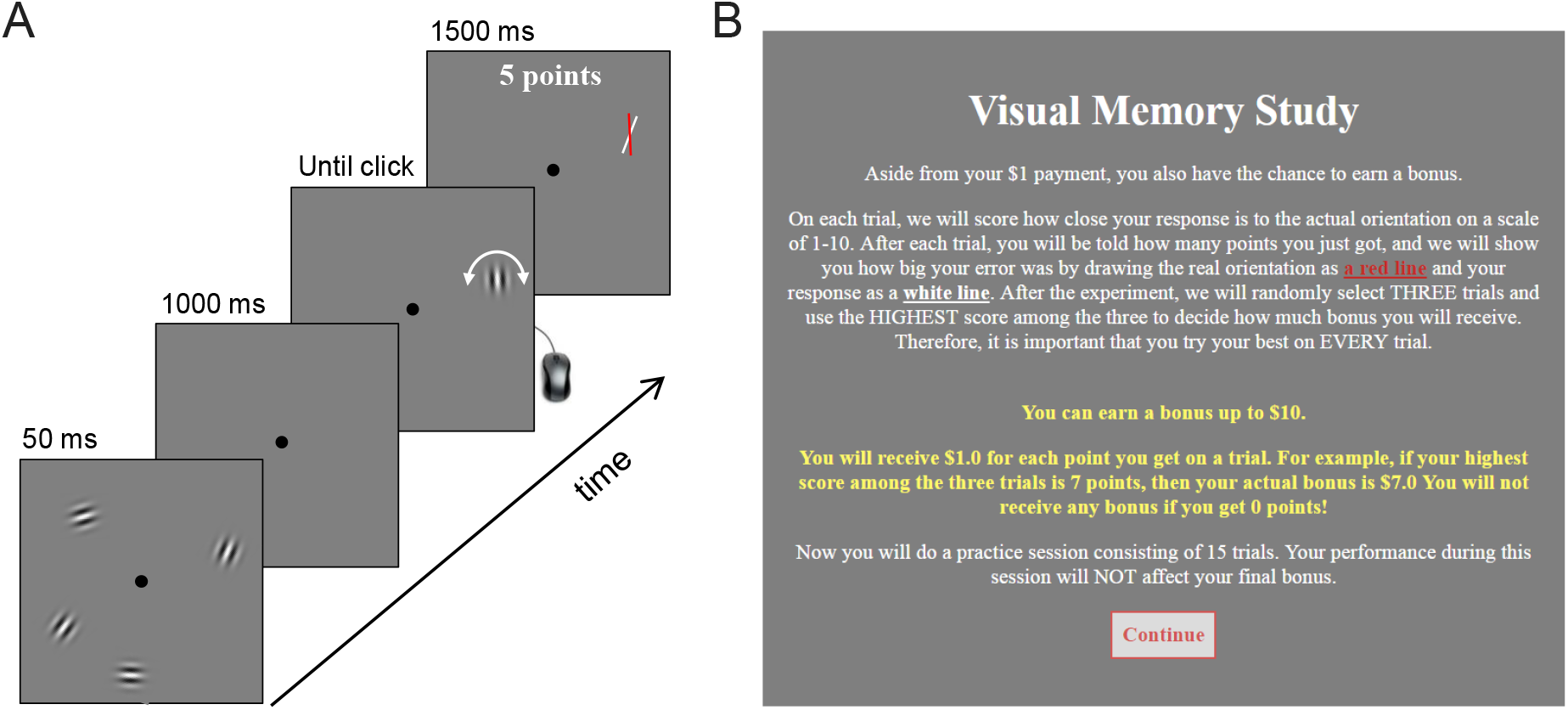
Experimental procedure. (A) Illustration of a single trial in Experiment 1 (not to scale). Subjects were briefly presented with 1,2,4, 6, or 8 Gabor patches, which they had to keep in memory during the delay period. Thereafter, a randomly oriented Gabor patch would appear at one of the previous stimulus locations. The task was to match the orientation of this stimulus with the remembered orientation of the stimulus that had appeared earlier at this location. The procedure in Experiment 2 was the same, except that no feedback was shown. (B) Instructions provided to the subjects in Experiment 1.

### Procedure

At the start of the experiment, subjects received written instructions about the task and about how their performance would be scored (Figure 1B). Next, they were informed about the bonus payment. For a subject in the condition with a maximum bonus of $10, the text in this screen would read “*After the experiment, we will randomly select THREE trials and use the HIGHEST score among the three to decide how much bonus you will receive […] You will receive $1 for each point you get on a trial. For example, if your highest score among the three trials is 7 points, then your actual bonus is $7. You will not receive any bonus if you get 0 points!”.* Thereafter, they performed 15 practice trials that were identical to trials in the actual experiment. After finishing these trials, a multiple-choice quiz was presented with three questions to test the subject’s understanding of the task and the potential bonus payment. Subjects who failed on at least one of these questions were disqualified from the experiment. The remaining subjects performed 250 trials of the delayed-estimated task with the five set sizes pseudo-randomly intermixed. To check if subjects were paying attention, we asked them at three points in the experiment to press the space bar within 4 seconds (catch trials). Subjects who at least once failed to do this were presumably not paying attention and were therefore excluded from the analyses.

### Results

Data from 10 subjects were excluded from the analyses because they failed to respond to at least one of the three catch trials. Of the remaining 189 subjects, another 35 were excluded because they had response error distributions that did not significantly differ from a uniform distribution, as assessed by a Kolmogorov-Smirnov test with a significance level of 0.05. The final group on which we performed the analysis thus consisted of 154 subjects, with 37, 41, 38, and 38 in the $0, $2, $6, and $10 conditions, respectively. To test for effects of reward level on VWM performance, we computed for each subject the circular variance^1^ of the response error distribution at each set size (Figure 2A, left). We performed a Bayesian Repeated-Measures ANOVA (JASP Team, 2018; Rouder et al., 2012) on these measures, with set size as a within-subjects factor and bonus level as a between-subjects factor. The results indicated extremely strong evidence for a main effect of set size (BF_incl_=∞), but evidence *against* a main effect of bonus level (BF_excl_=21)^2^.

**Figure 2.**
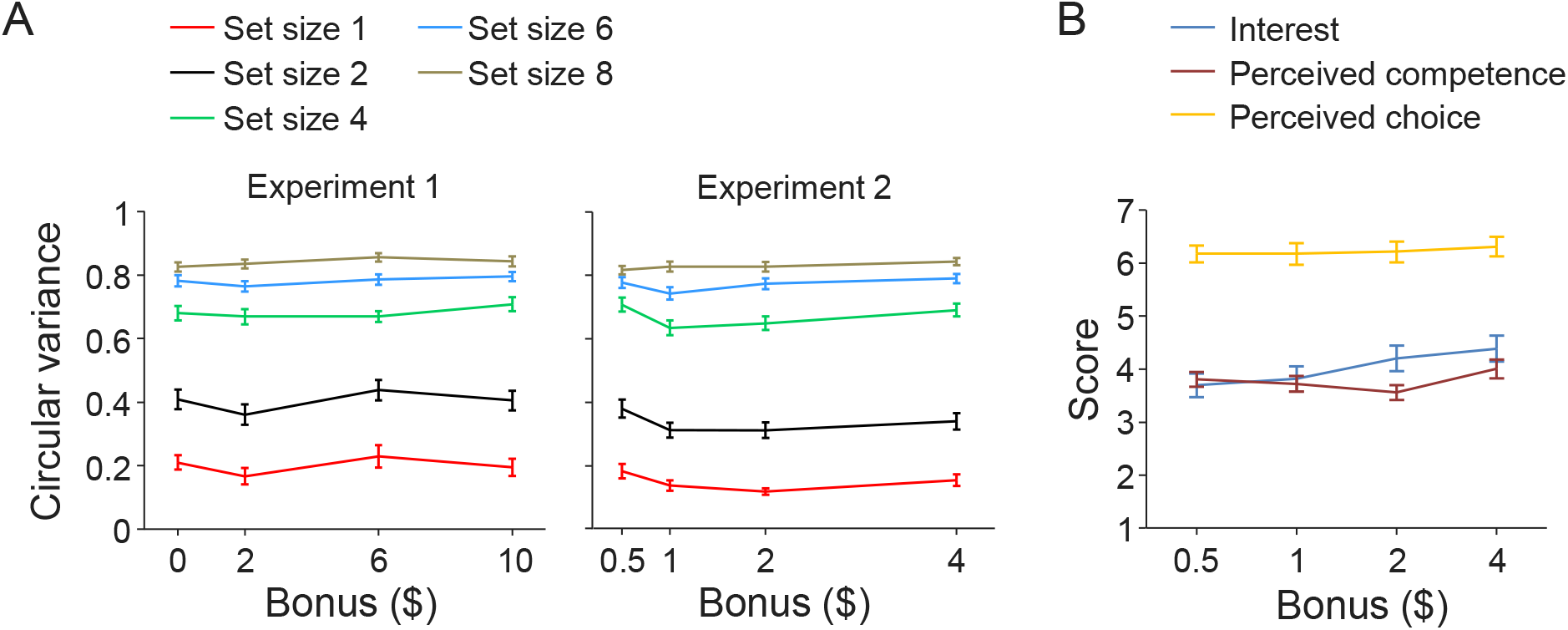
Effect of monetary reward level on memory precision and self-reported motivation scores in Experiments 1 and 2. (A) Subject-averaged circular variance of the estimation error distribution as a function of the amount of potential bonus in Experiments 1 (left) and 2 (right). (B) Intrinsic Motivation Inventory scores as a function of the amount of potential bonus, split by item category. Error bars indicate 1 s.e.m.

### Discussion

The results of Experiment 1 showed no evidence of an effect of performance-contingent reward on VWM performance. One possible explanation of this null result is that resource allocation in VWM is insensitive to monetary reward. However, there are at least two factors in the experimental design that may have interfered with the reward manipulation. First, subjects received trial-to-trial feedback. Being constantly confronted with their own performance may have motivated them to perform as well as possible regardless of the amount of bonus they could earn. Second, since the bonus was mentioned only at the beginning of the experiment, subjects may have performed the task without having the bonus strongly on their minds. To address these potential confounds, we ran a second experiment in which subjects did not receive trial-to-trial feedback and were reminded regularly of the bonus.

## EXPERIMENT 2

### Recruitment

A new cohort of subjects was recruited on the Amazon Mechanical Turk platform. The experiment was visible only to subjects who were located in the USA, had not participated in the experiment before, and had an approval rate of 95% or higher. A total of 241 subjects signed up, of whom 41 were disqualified due to failing the post-instruction quiz. The remaining 200 subjects were randomly assigned to four groups (*n*=52, 48, 50, 50) that again differed in the amount of potential bonus payment. The base payment was $5 and the potential bonus amounts were $0.50, $1, $2, and $4. The experiment was approved by the Institutional Review Board of New York University.

### Stimuli and procedure

The stimuli and procedure for Experiment 2 were identical to Experiment 1, except for the following differences. First, subjects were reminded of the bonus four times in the instruction screen (compared to only once in Experiment 1) and during the task itself the following message appeared after every 50 trials: *“You have completed X% of the Experiment. Remember that you have the chance to earn a $Y bonus!”,* where *X* and *Y* were determined by the number of completed trials and the amount of bonus, respectively. Second, no performance feedback was given, neither during practice nor during the actual experiment. Third, the length of the practice phase was reduced to 10 trials, but three “walk-through trials” were added at the start in which subjects were fully guided with additional written instructions. Lastly, after the experiment, subjects rated 20 items that we had selected from the Intrinsic Motivation Inventory (McAuley et al., 1989; Ryan, 1982). We only included the subscales “Interest/Enjoyment” (7 items, such as “This activity was fun to do”), “Perceived competence” (6 items, such as “I was pretty skilled at this activity”), and “Perceived Choice” (7 items, such as “I did this activity because I wanted to”) in our questionnaire. Subjects rated these items on a Likert scale from 1 (“not at all true”) to 7 (“very true”). An overview of all items is found in the Appendix.

### Results

Data from 27 subjects were excluded because they failed to respond to one of the catch trials (9 subjects) or had a response error distribution that did not significantly differ from a uniform distribution according to a Kolmogorov-Smirnov test (18 subjects). After these exclusions, we had 47, 42, 44, and 40 subjects in $0.50, $1, $2, and $4 conditions, respectively. We performed the same statistical analyses as in Experiment 1 on the data from the remaining 173 subjects (Figure 2A, right). Again, we found extremely strong evidence for a main effect of set size (BF_incl_=∞) and evidence *against* a main effect of bonus level (BF_excl_=2.9). Hence, it seems unlikely that the absence of an effect in Experiment 1 was due to subjects being unaware of the potential bonus payment or due to presence of trial-to-trial feedback.

Next, we assessed whether bonus size affected the subjects’ scores on the intrinsic motivation inventory questions (Figure 2B). Using Bayesian one-way ANOVAs, we found that there was no effect in any of the three categories: BF_10_=0.275 for mean “interest” scores, BF_10_=0.174 for mean “perceived competence” scores, and BF_10_=0.034 for mean “perceived choice” scores. Nevertheless, we noticed that there was considerable variation in the intrinsic motivation scores across subjects, especially in the “Interest” and “Perceived competence” categories (Figure 3A). Therefore, we next tested if there was an effect of motivation scores on VWM performance. To this end, we grouped subjects from Experiment 2 into “low motivation” and “high motivation” subgroups by using a median split on each of the three categories of the Intrinsic Motivation Inventory (Figure 3B). To examine whether scores in any of the three categories is predictive of VWM performance, we performed a repeated-measures Bayesian ANOVA with set size as within-subjects factor and motivation score (“low” and “high”) as a between-subjects factor. All three tests provided evidence for the null hypothesis that there was no performance difference between subjects in the low and high motivation subgroups (Interest: BF_excl_=8.3; Perceived competence: BF_excl_=4.8; Perceived choice: BF_excl_=4.8).

**Figure 3.**
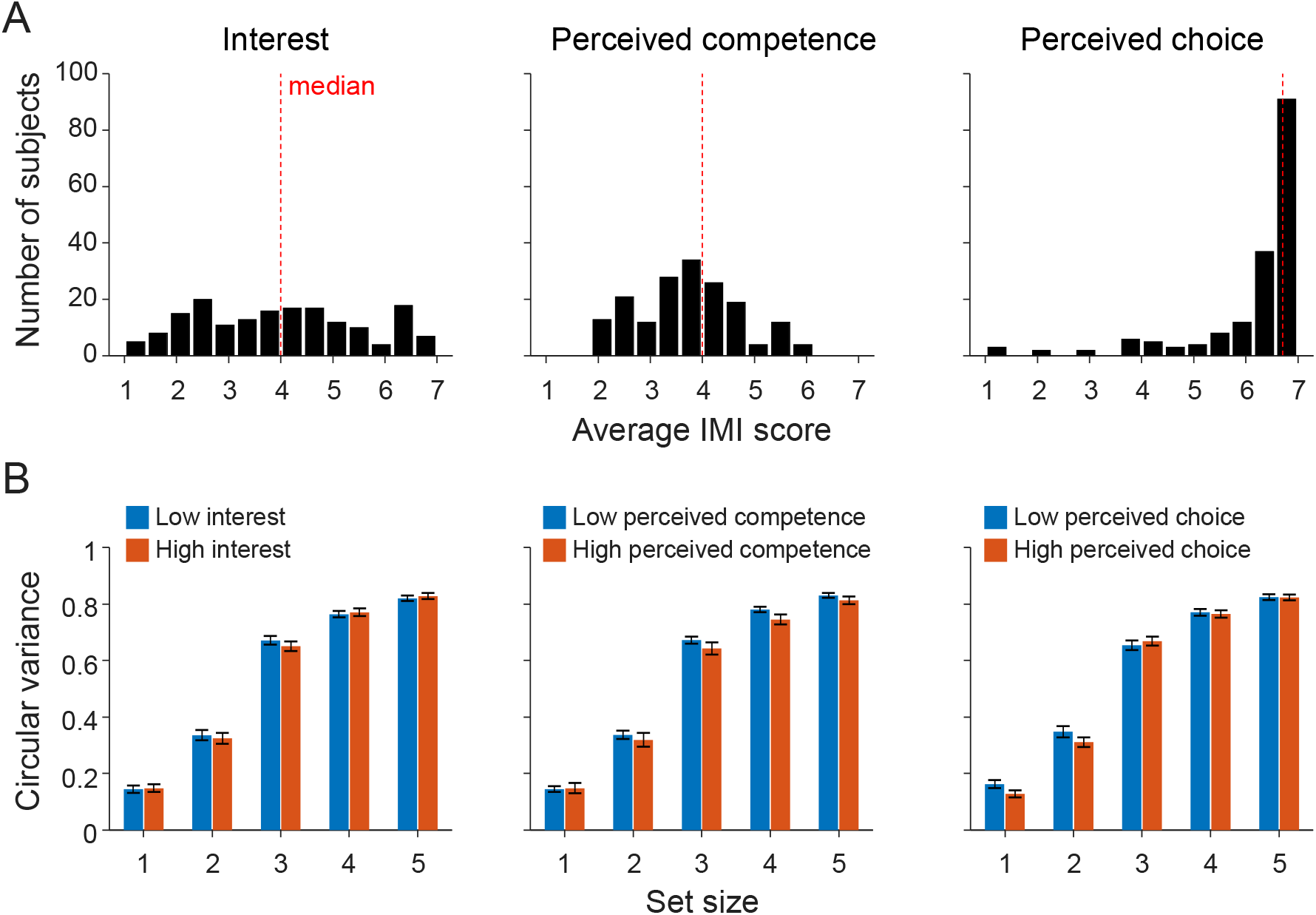
Comparison of VWM performance between subjects with low and high scores on the Intrinsic Motivation Inventory (IMI). (A) Distribution of average IMI scores, split by question category. (B) Circular variance of the response error plotted separately for subjects with below-median and above-median scores on the IMI questionnaire.

### Discussion

The aim of Experiment 2 was to test whether the null effect from Experiment 1 persists if we remove trial-by-trial feedback and remind subjects more often of the potential bonus. We found that this was the case: again, there was no effect of monetary reward on VWM performance. This further strengthens the hypotheses that VWM resource allocation is independent of monetary reward. We also found that intrinsic motivation did not depend on the amount of monetary reward. Therefore, our present results are limited to the realm of external motivation manipulations. It would be useful to also test for effects in a design where intrinsic motivation is experimentally manipulated.

In both Experiment 1 and Experiment 2, we manipulated reward using a between-subjects design. Hence, for each subject the expected reward was the same for each item on each trial. However, there are previous indications that effects of monetary reward on cognitive performance are *relative* rather than absolute, which can be detected using within-subject designs in which reward is manipulated trial-by-trial (e.g. Chiew & Braver, 2016; Engelmann et al., 2009; Etzel et al., 2016; Hall-McMaster et al., 2019; Kleinsorge & Rinkenauer, 2012; Locke & Braver, 2008; Poh et al., 2019; Shen & Chun, 2011). To test whether VWM encoding quality is affected by relative reward levels, we performed a third experiment in which reward was varied on a trial-by-trial level.

## EXPERIMENT 3

### Preregistration

We preregistered the methods of Experiment 3 at the Open Science Foundation platform (URL: https://osf.io/dkzax). The report below is in accordance with the preregistration, except when explicitly indicated otherwise.

### Recruitment

A third cohort of subjects was recruited on the Amazon Mechanical Turk platform. The experiment was visible only to subjects who were located in the USA, had not participated in the experiment before, and had an approval rate of 95% or higher. Data was collected from a total of 201 subjects, which were randomly assigned to two groups that differed in how the final payment was calculated (see below for details).

### Stimuli and procedure

Experiment 3 was similar to Experiments 1 and 2, but with a few important differences. Just as in the first two experiments, the instructions at the start of the experiment informed subjects that they would score points on each trial and that their score would be converted into a monetary bonus. Subjects received a score between 0 and 10 points on each trial, depending on how close their response was to the correct response. The mapping between the absolute error, *ε* (in degrees), and the score, *s*, was similar as in Experiments 1 and 2 (see Appendix). In the instructions, this mapping was visualized to the subject by showing an example error corresponding to each of the 11 possible scores, similar to the way feedback was provided in Experiment 1 (Figure 1A). Subjects were also informed that each trial had a *score multiplier* that could take values 1, 2, and 4. The multiplier was visualized as a text “1x”, “2x”, or “4x” at the center of the screen. To ensure that subjects would not forget about the multiplier, the text stayed on the screen throughout the entire trial (it also served as a fixation marker). The subject’s response score was on each trial multiplied by the trial’s score multiplier. At the end of the trial, the error was visualized using two lines (Figure 1A) and feedback about the gained score was provided by showing “[multiplier] x [response score] = [multiplied score]” at the fixation location (e.g., “2 x 6 = 12”).

After reading the instructions, subjects were presented with three walk-through trials (one for each multiplier). These self-paced trials explained once again the relation between the response error and the score as well as the role of the score multiplier. Next, they performed 18 practice trials that were identical to the experimental trials, but which were not included in the analyses. Thereafter, they were presented with three multiple-choice questions to check if they had understood the task and the way that their payment was calculated. Subjects who failed to answer all three questions correctly, were disqualified from performing the experiment. Subjects who passed the quiz would next start the main experiment, which consisted of 40 trials for each combination of set size (1, 3, or 6) and multiplier (1x, 2x, 4x), presented in random order (360 trials in total). Note that here we reduced the number of trials from 50 (in the preregistration) to 40, which was done to keep the experiment duration to approximately an hour. After performing the experiment, subjects were asked to indicate on a scale from 1 to 5 how much effort they you put in trials with a 1x, 2x, and 4x score multiplier (they provided three ratings, one for each multiplier).

### Two payment conditions (between-subject factor)

Since we measure hundreds of trials per subject, the expected reward per trial is quite low. A risk of this is that at the level of a single trial, subjects may perceive the expected bonus as so low that it is not really different from getting no bonus at all. Therefore, we divided subjects into two groups which differed in how their monetary bonus was calculated. In the first group, the bonus was calculated based on the summed score across *all* trials, with every 8 points being worth 1 cent. In the second group, the bonus was based on the total score of 9 trials (3 for each multiplier) that would randomly be selected after the experiment, with each 20 points being worth $1. These numbers were chosen such that the maximum total reward as well as the expected reward under guessing were the same in both groups. However, subjects in the second group may have felt that there was more at stake on any given trial than subjects in the first group (“what if this is one of the bonus trials?”). Note that the bonus calculations differed slightly from how we had formulated them in the preregistration (“Every 15 points in your total multiplied scores is worth 2 cents” for the all-trials group and “Every 15 points in your total multiplied score is worth $1” for the nine-random-trials group), but the underlying idea was still the same.

### Results

A total of 52 subjects were excluded because they failed to respond to one or more of the catch trials. Another 49 subjects were excluded because a Kolmogorov-Smirnov test failed to reject the hypothesis (at a *p* < 0.05 level) that their response error distribution was uniform, suggesting that they submitted random responses. The analyses reported below were performed on the 100 remaining subjects (with 52 and 48 in the first and second payment group, respectively). Unfortunately, due to a technical error, the post-experiment questionnaire data were collected for only a few of the subjects; we did not use these data in any of our analyses.

We performed a Bayesian ANOVA with performance (measured as the circular variance of the response error) as the dependent variable, multiplier and set size as within-subject factors, and payment condition as a between-subject factor. The results revealed extremely strong evidence for a set size effect (BF_incl_ > 10^13^), but no evidence for an effect of payment condition (BF_excl_ = 9.0) or multiplier (BF_excl_ = 38) (Figure 4).

**Figure 4.**
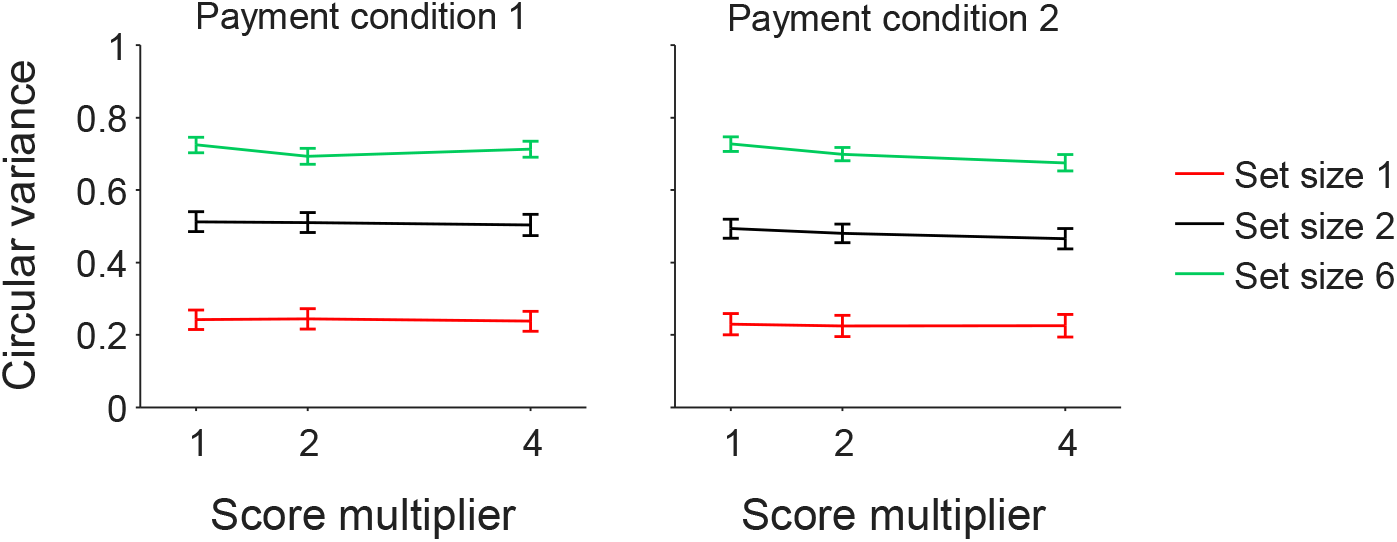
Effect of monetary reward level on memory precision in Experiment 3. (A) Subject-averaged circular variance of the estimation error distribution as a function of the trial-by-trial score multiplier. Payment Condition 1 refers to the group of subjects whose bonus was calculated on the score summed across all trials and Payment Condition 2 to the group whose bonus was calculated on 9 randomly selected trials. Error bars indicate 1 s.e.m.

While not specified in the preregistration, we also performed a linear mixed modelling analysis using the *lme4* package for R (Bates et al., 2015). This analysis is possibly more powerful in detecting effects, because it can deal with individual differences in baseline performance. Set size, multiplier, and payment condition were specified as fixed effects and the intercept for subjects as a random effect. To estimate the evidence for an effect, we performed a Chi-square test between the full model and the model with the effect removed. Consistent with the ANOVA, this provided strong evidence for an effect of set size (p < 0.0001), but no evidence for an effect of multiplier (*p*=0.14) or payment condition (*p*=0.56).

Next, we analyzed the data at the level of individuals by performing the model-based analysis as planned in the preregistration. For each subject, we first fitted a variable-precision model (van den Berg et al., 2014, 2012) in which mean encoding precision, 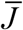, depended only on set size, *N*, through the power-law relation 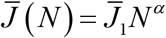, where parameter 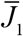 determined the precision at set size 1 and parameter *α* how fast precision decreases with set size, respectively. We compared the goodness of fit of this model to a variant in which encoding precision also depended on the reward level, by fitting 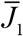 separately for trials with multipliers 1, 2, and 4 (thus introducing two additional parameters). The second model has an AIC value that is on average only 3.11 ± 0.21 points lower than that of the first model, which indicates that the models describe the data approximately equally well. In other words, the model comparison result is consistent with the above statistical analyses and indicates that there is no evidence for an effect of reward level on encoding precision.

### Discussion

As in the first two experiments, we did not find any indications for an effect of monetary reward level on VWM performance. The main difference with the first two experiments is that Experiment 3 manipulated relative rather than absolute reward, by varying the potential monetary bonus from trial to trial. Another new manipulation in this experiment was the way in which bonus was calculated: for half the subjects the bonus was based on *all* trials while for the other half is was based on 9 randomly selected trials. We found no effect of this manipulation either on VWM precision, which makes it less likely that the null effects are due to subjects perceiving the per-trial bonus as too low to have any effect on their motivation.

## CONCLUSION

In three experiments, we found no evidence that encoding precision in VWM depends on performance-contingent monetary reward. One could argue that our null effects may have been the result of using a “flawed” experimental design and that we would have found effects if we had done the “right” experiment. Even so, the fact that we found no effect in three different experiments that varied a large number of experimental factors would limit the generality of any relation that may exist between monetary reward and working memory performance; if money has an effect on VWM precision, quite specific conditions seem to be required to detect it. We consider multiple explanations for the apparent dissociation between monetary reward and VWM performance.

First, the bonuses may have been too small to cause an effect. We believe this to be unlikely, especially in Experiment 1, where the bonus could increase the earnings in one of the groups by a factor 11 ($10 bonus in addition to $1 base payment).

Second, subjects might not have had the bonus payments strongly enough on their minds when performing the task. While this explanation could be plausible in Experiment 1 – where subjects were informed about the bonus only at the very beginning of the experiment – it seems implausible in Experiments 2 and 3, where they were regularly reminded of it.

Third, we may inadvertently have biased our subject sample to “over-performers”, by only recruiting subjects who had a high approval rate on the Amazon Turk. The desire to maintain a high approval rate may have worked as a strong incentive for these subjects to perform well, regardless of the amount of performance-related bonus they could earn. However, we know of at least two other studies that have found effects of monetary reward on performance in Amazon Turk subjects with high approval rates, albeit using non-memory tasks (Caplin et al., 2020; Dellavigna & Pope, 2018).

Fourth, the present study only manipulated *external* reward. An interesting direction for further study would be to test for effects of *intrinsic* motivation on VWM performance, for example by “gamifying” the experiment - using concepts such as leaderboards, achievements, and levels – which have shown to have positive effects in other contexts such as education and learning (Hamari et al., 2014).

Finally, it may be that VWM uses a fixed amount of resource, independent of the task at hand. This explanation is consistent with another recent study that also found no effect of reward on total capacity in an orientation estimation task similar to ours (Brissenden et al., 2021). However, it is inconsistent with yet another recent study that did find an effect, albeit with modest effect sizes and only in one of their two subject groups (Manga et al., 2020). Moreover, the idea of a fixed capacity contradicts previous evidence suggesting that the amount of allocated resource depends on task instructions (Bengson & Luck, 2016), set size (van den Berg & Ma, 2018), and cueing condition (Myers et al., 2018). Moreover, this kind of rigidity would stand in stark contrast to the flexibility with which VWM resource is divided among items within a trial when items have varying importance (Bays, 2014; Bays et al., 2011; Emrich et al., 2017; Gorgoraptis et al., 2011; Yoo et al., 2018; Zokaei et al., 2011). Altogether, VWM seems to be a flexible system that can adjust its capacity to certain external factors, but monetary reward does not may not be one of them.

## ACKNOWLEDGMENTS

This research was supported by grant 2018-01947 from the Swedish Research Council to R.v.d.B, training grant R90DA043849-03 to Q.Z., and grant R01EY020958-09 to W.J.M.

## APPENDIX Model predictions

Figure A1 presents results from a simulation analysis using our earlier proposed resource-rational model of visual working memory (van den Berg & Ma, 2018), applied to a single-probe delayed-estimation task (Blake et al., 1997; Prinzmetal et al., 1998; Wilken & Ma, 2004). The results reveal that the model predicts strong effects of both set size and reward level on the circular variance of the estimation error, as well as an interaction effect.

**Figure A1.**
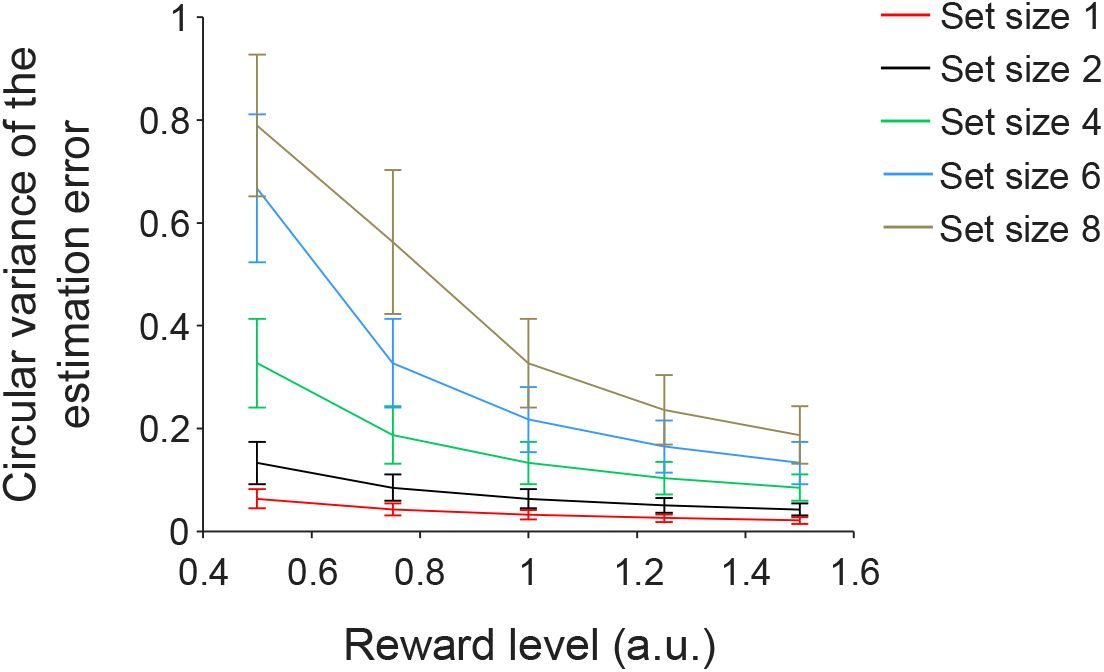
Effect of reward on the circular variance of the estimation error in a delayed-estimation task, as predicted by a resource-rational model of visual working memory. The predictions were obtained by simulating responses of the model presented in (Van den Berg & Ma, 2018). Simulations were performed at five set sizes (separate lines) and 5 reward levels (*x*-axis). Each of the simulations was performed six times, with run using the maximum-likelihood parameters of one of the six subjects in experiment E4 of that paper (which used an orientation task, as in the present paper). Error bars represent ±1 SEM across the six runs. A two-way Bayesian ANOVA on the simulation data show strong evidence for an effect of both set size (BF_incl_ > 10^13^) and reward level (BF_incl_ > 10^8^), as well as for an interaction effect (BF_incl_ = 98).

### Scoring functions

In all three experiments, subjects received points on each trial based on the accuracy of their estimate. In Experiment 1, errors were mapped to scores through the function 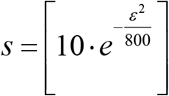, where *ε* is the error in degrees and [.] indicates rounding to the nearest integer (Fig A1, black). In Experiments 2 and 3, the function were 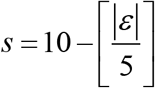 and 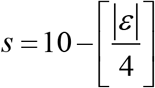, respectively, giving highly similar mappings as in Experiment 1 (Figure A1, red and green).

**Figure A2.**
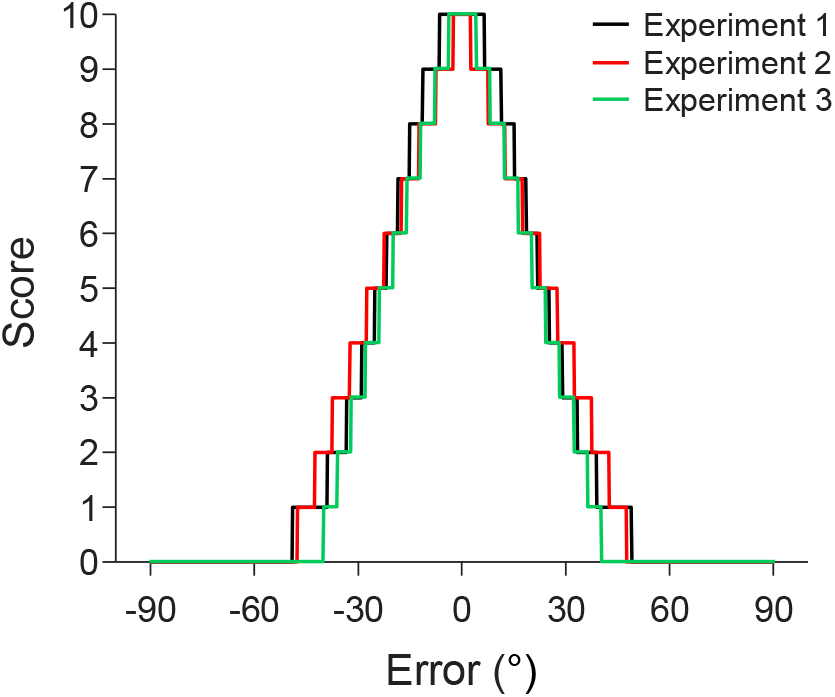
Scoring functions used in the three experiments.

### Questionnaire items in Experiment 2

Subjects in Experiment 2 filled out a questionnaire with the following items from the Intrinsic Motivation Inventory (McAuley et al., 1989; Ryan, 1982):

- Interest/Enjoyment

- I enjoyed doing this activity very much
- This activity was fun to do.
- I thought this was a boring activity. (R)
- This activity did not hold my attention at all. (R)
- I would describe this activity as very interesting.
- I thought this activity was quite enjoyable.
- While I was doing this activity, I was thinking about how much I enjoyed it.
- Perceived Competence

- I think I am pretty good at this activity.
- I think I did pretty well at this activity, compared to other students.
- After working at this activity for a while, I felt pretty competent.
- I am satisfied with my performance at this task.
- I was pretty skilled at this activity.
- This was an activity that I couldn’t do very well. (R)
- Perceived Choice

- I believe I had some choice about doing this activity.
- I felt like it was not my own choice to do this task. (R)
- I didn’t really have a choice about doing this task. (R)
- I felt like I had to do this. (R)
- I did this activity because I had no choice. (R)
- I did this activity because I wanted to.
- I did this activity because I had to. (R)

Subjects rated these items on a Likert scale from 1 to 7. Scores on items indicated with an (R) were reversed before entering them into the analysis.

## Footnotes

1 The circular variance was computed as 1 – *R*, where *R* is the length of the resultant vector of the subject’s estimation errors, measured as the circular distance between the true orientation and the response.

2 BF_incl_ measures how likely the data are in models that *include* the factor compared to how likely they are in models *exclude* the factor. Likewise, BF_excl_ (=1/BF_incl_) measures how likely the data are in models that *exclude* the factor compared to how likely they are in models that *include* the factor.

## Notes

https://osf.io/mwz27/

### Competing Interest Statement

The authors have declared no competing interest.

### Summary of Updates

We added a third experiment with a within-subject design to test if the lack of effects in Experiments 1 and 2 may have been due to only using between-subject reward manipulations.

